# Postmortem findings of free-ranging North American beavers (*Castor canadensis*) reveal potential threats to California freshwater ecosystems

**DOI:** 10.1101/2024.09.21.614286

**Authors:** Omar A. Gonzales-Viera, Leslie W. Woods, Aslı Mete, Heather Fritz, Anibal G. Armien, Emma Lantz, Luis A. Gomez-Puerta, Daniel Famini, Jaime Sherman, Jaime L. Rudd, Lauren E. Camp, Karen Shapiro, Deana L. Clifford

## Abstract

North American beavers (*Castor canadensis*) are semi-aquatic rodents recognized as keystone species that increase the diversity of freshwater ecosystems. This study aimed to characterize the mortality and pathological findings of free-ranging beavers in California and, based on these results, establish the potential threats to freshwater ecosystems. This study included eighteen beavers submitted for postmortem examination between 2008 and 2024 at the California Animal Health and Food Safety Laboratory, UC Davis. Gross and microscopic examination, bacteriological, parasitological, immunohistochemical, and molecular techniques were used as tools to diagnose the cause of death/reason for euthanasia and comorbidities in the beavers. Encephalitis caused by the larva migrans of *Baylisascaris* spp. was the most prevalent (9/18) cause of mortality/reason for euthanasia followed by bacterial infections in 6 individuals. In these 6 animals, bacterial bronchopneumonia was diagnosed in two (*Staphylococcus aureus* and a mix of gram-negative and -positive bacterial infection) and *Listeria monocytogenes* encephalitis, bacterial myofascitis (*Aeromonas bestiarum* and *Pasteurella multocida*), bacterial encephalitis (*Acinetobacter towneri*), and tularemia (*Francisella tularensis*) were diagnosed in one beaver each. Three animals died or were euthanized due to non-infectious causes including motor vehicle trauma, squamous cell carcinoma, and capture cardiomyopathy. Endoparasitism was the main comorbidity including granulomatous hepatitis by a suspected capillarid species, cerebral toxoplasmosis, Giardia infection, gastric nematodiasis, and cecal trematodiasis. In California, beavers are exposed to various pathogens that represent threats to humans, domestic animals, and wildlife. Since the interspecies transmission of these pathogens occurs in rivers, streams, lakes, and ponds, we suggest that studying beaver health can reflect freshwater ecosystem health. This study also indicates that the translocation of beavers into new areas without consideration and/or mitigation represents a potential risk for pathogen introduction.

## Introduction

Freshwater ecosystems such as lakes, reservoirs, rivers, streams, and wetlands are interactive systems with unique dynamics and diversity [1]. These water systems provide an aquatic or semi-aquatic environment that can support native fauna and flora [2,3]. Unfortunately, the inherent value of these ecosystems attracts human production and recreational activities, livestock husbandry, and invasive species causing detrimental consequences for freshwater health [4,5]. The health status of these biological communities can be evaluated by studying the water-borne infectious diseases, water pollutants, mosquito-borne pathogens, and diseases affecting the inhabitant native fauna.

Animals are an important sentinel of the health of aquatic freshwater systems since they are continuously exposed to multiple environmental stressors [6,7] that might result in diseases and mortality. North American beavers (*Castor canadensis*) are herbivores and semi-aquatic mammals adapted to live in and around freshwater sources [8]. Beavers are considered “ecosystem engineers” because of their ability to modify ecosystems by building dams or “keystone species” due to their fundamental role in the survival of multiple species in a landscape [8,9,10]. Beavers spend a large portion of their lifespan in freshwater systems making them prone to threats from water-borne infectious diseases and impacts of anthropogenic activities [3,7,9]. Numerous bacteria with zoonotic potential such as *Francisella tularensis* (type B, typically linked to an aquatic cycle) [11] or *Yersinia pseudotuberculosis* and *Y. enterocolitica* have caused mortality of free-ranging beavers [12,13]. Parasites such as *Cryptosporidium* spp., *Giardia* spp. [3,14,15], *Toxoplasma gondii*, and *Sarcocystis neurona* [15] are among the main water-borne protozoans infecting beavers in North America, while the neural larva migrans (NLM) of *Baylisascaris* spp. has caused encephalitis in beavers and other rodents sharing the same niche with raccoons [16]. Rabies virus with presumptive pantropism was diagnosed in a free-ranging American beaver that bit a man in Pennsylvania, USA [17]. In addition to infectious diseases, hazardous concentrations of pollutants such as cadmium, mercury, and other metals have been reported in beavers from different regions [7,18,19]. Since most of the studies regarding North American beaver mortality and pathology are sparse and the majority involve brief reports, we compiled a set of cases from California focusing predominantly on infectious diseases. Our findings suggest that beaver health conditions can serve as good indicators of the water quality therefore freshwater ecosystem health in California.

This study aims to characterize the mortality and pathological findings of free-ranging North American beavers submitted to the California Animal Health and Food Safety (CAHFS) Laboratory system by the California Department of Fish and Wildlife (CDFW) or rehabilitation centers in California with a special focus on infectious diseases.

## Materials and Methods

### Study design

This study combined retrospective and prospective cases submitted for postmortem diagnostic investigation to the CAHFS Laboratory system, where the beaver cadavers were examined by American College of Veterinary Pathologists board-certified anatomic pathologists. For the retrospective study, data (e.g., clinical history, necropsy reports, and ancillary diagnostic tests) of all free-ranging North American beavers submitted to the lab from January 2008 to December 2018 were retrieved from the Laboratory Information Management System digital platform and carefully evaluated. Furthermore, glass slides stained with hematoxylin & eosin (H&E) were retrieved from the archives to be re-evaluated. The prospective study involved all North American beavers submitted to CAHFS between January 2019 and May 2024, which were examined by the first author (OAGV) along with three other pathologists (AM, AGA, and LWW). Formalin-fixed paraffin-embedded (FFPE) blocks from these cases were selectively used when recuts, immunohistochemistry or special stains were necessary. Virological, bacteriological, parasitological, and molecular tests were requested upon consideration of the pathologist after gross or microscopic examination of each case.

### Demographic data and clinical information

Sex, age class (adult or juvenile), date and location found, and manner of death (natural death or euthanasia) were recorded for all collected beavers. Cadavers were submitted for postmortem examination along with summarized clinical information written by a wildlife veterinarian (supporting information S1 Table1).

### Postmortem examination

Gross examination and tissue sampling were performed following standard protocols for necropsy of mammals.

### Histology

Formalin-fixed tissues were routinely processed and cut at 4-µm sections for H&E staining. Gram, Periodic Acid-Schiff (PAS), Gomori’s methenamine silver (GMS), Masson’s trichrome, Congo red, Perl’s, and acid-fast stains were performed following the standard operating procedures (SOP) of the histology laboratory, CAHFS, Davis Lab.

### Immunohistochemistry

Four-micrometer sections of FFPE tissues were used for immunohistochemistry to Pancytokeratin (PanCK), vimentin, *Listeria monocytogenes*, *Toxoplasma gondii*, *Neospora caninum*, *Sarcocystis neurona*, West Nile virus, canine distemper virus (CDV), and *Francisella tularensis* following specific SOPs of the histology laboratory, CAHFS, Davis Lab. Procedures included 3% hydrogen peroxide treatment in water for 10 min after deparaffinization and rehydration of the tissue sections. Specific details of the use of the primary antibodies and positive controls are shown in the supporting information (S2 Table). The primary antibody was diluted in DaVinci Green antibody diluent (PD900M, Biocare Medical) and incubated for 45 min at room temperature. The secondary antibodies (Dako, Envision System-HRP) anti-mouse or anti-rabbit were incubated for 30 min at room temperature. Tris-buffered saline-Tween was used for rinses between steps. Slides were visualized with 3-amino-9-ethylcarbazole chromogen (Ready-to-Use, K3464, Dako, CA), and counterstained by Mayer’s Hematoxylin. Negative control slides were included using the same technique but changing the primary antibodies with an unrelated antibody with the same isotype. Positive controls are displayed in S2 Table.

### Virology

Sections of the hippocampus, brainstem, and cerebellum of all but two of the beavers submitted with or without a history of neurological signs (ataxia, nystagmus, torticollis, or swimming in circles) were collected for immunofluorescence for rabies virus and processed according to the SOPs of the California Department of Public Health (CDPH), Richmond, CA.

### Bacteriology

Swabs were utilized to sample tissues during necropsy and transferred to the bacteriology laboratory at CAHFS in transport media (BD BBL™ CultureSwab Collection and Transport System with Liquid Stuart Medium). For routine aerobic cultures, 5% sheep blood agar (SBA), MacConkey agar (MAC), and chocolate agar (CHOC) were inoculated with a swab and streaked for isolation. Plates were incubated at 37°C +/- 2°C with 5-10% CO_2_ and evaluated at 24 and 48 hours for growth.

For specific recovery of *Listeria monocytogenes*, a small piece of tissue (or tissue swab) was transferred into Brain Heart Infusion (BHI) broth for cold enrichment by incubating at a refrigerated temperature (4°C) for 4 weeks. Once a week, the cold enrichment broth was subcultured onto SBA and Oxford agar (OX) plates and incubated at 37°C +/- 2°C with 5-10% CO_2_, with evaluation for growth at 24 and 48 hours. Bacterial isolates were identified using a combination of biochemical testing and matrix-assisted, laser desorption-ionization time of flight (MALDI-TOF) mass spectrometry (Bruker, MA).

To detect *Salmonella* spp., swabs were transferred into 10 mL of Tetrathionate (TT) broth supplemented with 0.2 mL Iodine and 0.1 mL 0.1% Brilliant Green solution and incubated at 37°C +/- 2°C for 18-24 hours. Following incubation, an aliquot of the TT-enriched sample was removed for DNA extraction, and Real-Time PCR (RT-PCR) was done using the IDEXX Real PCR reagents and an ABI 7500 Fast Real-Time PCR System to detect *Salmonella*. If the sample was positive by RT-PCR, it was subcultured to XLT-4, BGN, 5% SBA, and MAC plates and incubated at 37°C +/- 2°C and evaluated at 24 and 48 hours for suspect *Salmonella* colonies. *Salmonella* was identified using a combination of MALDI-TOF and biochemical tests. To specifically culture for *Yersinia* spp. from feces, a swab was collected for direct culture onto Cefsulodin-Irgasan-Novobiocin (CIN) selective media and MAC plates and incubated aerobically at 24-26°C for 42-48 hours, observing plates at 18-24 hours and 42-48 hours. Suspect colonies were identified by MALDI-TOF. In addition to the direct culture of samples, 0.5-1 gram of feces was transferred into a tube containing 5 mL sterile phosphate-buffered saline (PBS) and refrigerated at 3-5°C for 3 weeks. The sample in saline was subcultured to CIN and MAC plates weekly for 3 weeks and evaluated for suspect colonies with direct culture methods.

Samples from animals suspected to have tularemia were submitted to the CDPH and were cultured according to their SOPs.

### Parasitology

Fecal flotation was performed by double-centrifugation using sodium nitrate (Fecasol) flotation solution (specific gravity 1.20-1.27) to float ova onto a coverslip. Coverslips were transferred to a glass slide and parasite ova were visualized by light microscopy with the 10X and/or 20X objective. Types of ova and numbers detected were recorded.

Nematodes were clarified in an ethanol-phenol solution (1:2 v/v) to improve visualization of the organs. The anterior and posterior regions of the nematodes were examined, focusing mainly on the males. Trematodes were stained with hydrochloric carmine and Gomori’s trichrome, then dehydrated in successive ethanol series until absolute ethanol. Finally, they were clarified in eugenol and mounted on slides using Canada balsam. Parasites were observed under an optical microscope and a stereomicroscope. Nematode and trematode identification were performed using taxonomic keys [20,21,22].

### Molecular analysis

To confirm the presence of *T. gondii* infection in one beaver. DNA from the brain tissue was extracted in triplicate (DNeasy Blood and Tissue Kit (QIAGEN)) and amplified via nested Polymerase Chain Reaction (PCR) targeting the pan-apicomplexan first internal transcribed spacer 1 (ITS-1) locus [23]. An additional locus (B1) was amplified to characterize parasite genotype as previously described [24]. Briefly, the PCR reaction (50 μl volume) included 0.5 μl each of 50 μM forward and reverse primers, 0.8 μl of 10% Bovine Serum Albumin, 25 μl of 2X Amplitaq Gold 360MM, and 5 μl (external) or 2 μl (internal) DNA template. The PCR run contained one positive control (DNA derived from *T. gondii* ME49 (Type II) strain) and three negative controls (extraction reagents with distilled water, PCR reagent control, and another PCR reagent control with distilled water added in the nested reaction step). PCR amplification products were separated and visualized with gel electrophoresis on a 2% agarose gel stained with Red Safe and viewed with a UV transilluminator. Amplified bands were cut and purified from the gel using the Qiagen Qiaquick Gel Extraction kit and submitted for Sanger sequencing at the UCDNA Sequencing Facility. The forward and reverse sequences were aligned and trimmed using Geneious software (R11 Biomatters Ltd., Auckland, New Zealand), and the consensus sequence was compared with the GenBank database using the Basic Local Alignment Search Tool, BLAST (http://blast.ncbi.nlm.nih.gov/Blast.cgi). Parasite genotype at the B1 locus was first determined via virtual restriction fragment length polymorphism (RFLP) at the B1 locus using Geneious software. To identify a specific SNP location in sequences that did not align at 100% identity with previously reported sequences, we performed a local alignment in Geneious using common reference Types including Type I (ATCC RH strain), Type II (ATCC ME49 strain), Type III (ATCC CTG strain), Type X (KM243033 Bobcat strain), and an X variant strain (sea otter MK988572 strain).

Molecular testing was also conducted on frozen samples and on FFPE scrolls of brain and liver tissues to detect DNA of *Baylisascaris* spp. and capillarid nematodes on 7 and 1 beaver, respectively. For the frozen liver, DNA extraction proceeded using the same protocol described above. For FFPE scrolls, paraffin was first removed from the scrolls using a xylene substitute protocol. Briefly, nine 10-µm scrolls were processed in triplicate, with three scrolls each in three, 2-ml centrifuge tubes. Xylene substitute (1400 µl) was added to each of the tubes, which were vigorously vortexed and then centrifuged at 14,000 rpm for 3 minutes. After discarding the supernatant, 1400 µl of absolute ethanol was added to each tube followed by vortexing and centrifugation as above; a second ethanol wash was then performed in the same way. Following the second ethanol wash, the tubes were quickly centrifuged and any remaining supernatant was removed. DNA was extracted from the resulting pellets following the same procedure as for *T. gondii*. PCR was then conducted on these extracts using nematode-specific primers targeting the large subunit (28S) nuclear ribosomal DNA (rDNA) with previously established cycling conditions [25] and a *B. procyonis* positive control. PCR amplicons were purified and sequenced as above.

## Results

### Animals and clinical information

A total of 23 free-ranging beavers were submitted to CAHFS from January 2008 to May 2024. Four cadavers were excluded from the study due to advanced autolysis, and lack of adequate sample preservation and clinical history. A fifth beaver was excluded due to the excessive time spent in rehabilitation (>2 months) that might have influenced the encountered diseases. Thus, 18 beaver cadavers (9 females and 9 males) were included in this study. Twelve beavers were adults and 6 were juveniles. Twelve beavers were found dead while 6 were euthanized. Complete information on each animal is in Table 1. All the beavers inhabited Northern California and were clustered in highly populated counties from the Sacramento and San Francisco Bay areas (Figure 1). Contra Costa and Sacramento counties had the most submissions with 7 and 4 beavers, respectively.

**Figure 1.**
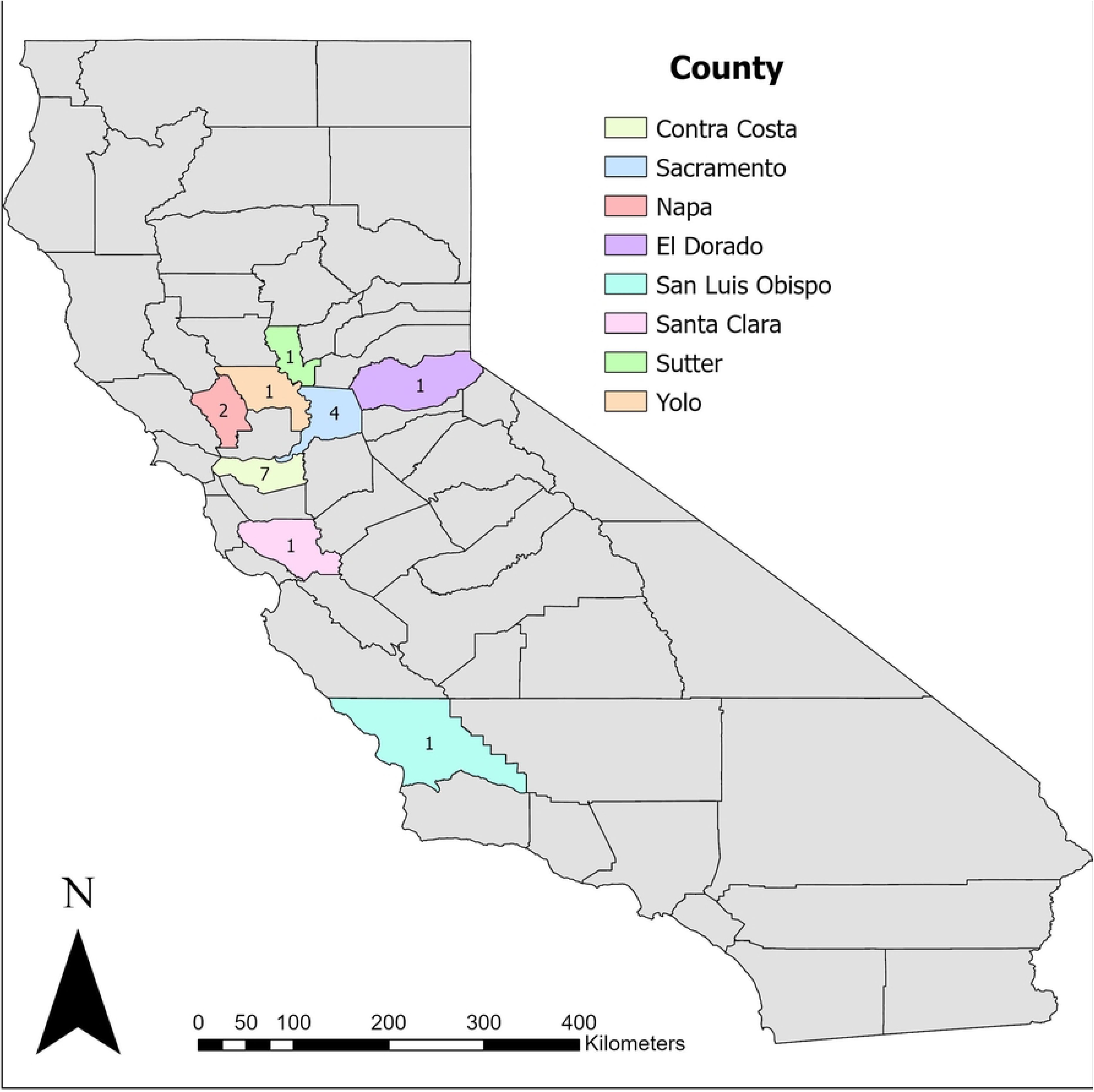
Geographic distribution at the county level of the studied beavers in California. The number of animals is shown within each county.

**Table 1.** Biological information, location found, cause of death or reason for euthanasia, and pathology of free-ranging beavers in California.

| Beaver | Sex | Age | Location | Cause of death or reason for euthanasia | Pathological findings |
| --- | --- | --- | --- | --- | --- |
| 1 | Male | Juvenile | Martinez, Contra Costa County | Baylisascariasis | Severe multifocal necrotizing and eosinophilic granulomatous encephalitis with intralesional neural larva migrans. Disseminated *Baylisascaris* sp. migration with granulomatous thymitis, pneumonia, myocarditis, hepatitis, glossitis, omentitis, enteritis, and diaphragmatic myositis |
| 2 | Female | Juvenile (1y) | Martinez, Contra Costa County | Baylisascariasis | Moderate multifocal glial scars with mild nonsuppurative meningoencephalitis and plexus choroiditis. Pulmonary edema and congestion with intra-alveolar hemorrhages. Mild interstitial nephritis, hepatitis, and myocarditis. |
| 3 | Female | Juvenile (1y) | Martinez, Contra Costa County | Baylisascariasis | Moderate multifocal glial scars with moderate non-suppurative encephalitis. Hepatocellular swelling with intracytoplasmic eosinophilic vacuoles. |
| 4 | Female | Adult (6y) | Martinez, Contra Costa County | Baylisascariasis | Multifocal glial scars. Eosinophilic and histiocytic interstitial pneumonia with vascular thrombi. Malocclusion with secondary ulcerative proliferative gingivitis. Lymphocytic polyneuritis. Lymphocytic and neutrophilic conjunctivitis. |
| 5 | Male | Juvenile (1y) | Pittsburg, Contra Costa County | Baylisascariasis | Severe multifocal eosinophilic granulomatous encephalitis with intralesional neural larva migrans, Disseminated *Baylisascaris* sp. migration with granulomatous lymphadenitis, myocarditis, enteritis and nephritis with visceral larva migrans. |
| 6 | Female | Adult | Winters, Yolo County | Baylisascariasis | Moderate multifocal glial scars. Moderate multifocal nonsuppurative, eosinophilic encephalitis. Disseminated *Baylisascaris* sp. migration with granulomatous hepatitis, lymphadenitis, omentitis, and splenitis with visceral larva migrans. Cholangiocellular adenoma. Papillary renal adenoma. Gastric nematodiasis. Cecal trematodiasis. |
| 7 | Female | Adult | Martinez, Contra Costa County | Baylisascariasis | Moderate multifocal glial scars and non-suppurative, eosinophilic encephalitis. Disseminated *Baylisascaris* sp. migration with granulomatous myocarditis, hepatitis, splenitis, gastritis, enteritis and colitis with visceral larva migrans. Moderate chronic interstitial pneumonia. Emaciation. Cecal trematodiasis. |
| 8 | Female | Adult | Palo Alto, Santa Clara County | Baylisascariasis | Severe multifocal granulomatous encephalitis with neural larva migrans. Severe multifocal glial scars. Severe diffuse chronic proliferative and lymphohistiocytic interstitial pneumonia with pneumocyte type II hyperplasia, fibrosis, smooth muscle hyperplasia, and rare plant-induced granulomas. Moderate multifocal fibrotic and papillary pleural hyperplasia. Mild multifocal lymphoplasmacytic tracheitis. Moderate multifocal fibrinonecrotizing colitis with mix-shaped bacteria |
| 9 | Female | Juvenile | Discovery Park, Sacramento County | Baylisascariasis | Severe multifocal eosinophilic granulomatous encephalitis with intralesional neural larva migrans, axonal degeneration, and multifocal glial scars. Disseminated *Baylisascaris* sp. migration with granulomatous pneumonia, enteritis, myocarditis, and lymphonodular capsulitis with visceral larva migrans. Skin, subcutis, and latissimus dorsi muscle (right side of the thorax): Severe focally extensive chronic active necrotizing dermatitis, cellulitis, and polyphasic myositis with mix-shaped bacteria. Subcutis and muscles (right scapula and humerus): Severe multifocal chronic active necrosuppurative cellulitis and myositis with mix-shaped bacteria. Severe diffuse acute superficial necrotizing colitis with mix-shaped bacteria. |
| 10 | Male | Adult | Lake Tahoe, El Dorado County | Tularemia | Severe acute random necrotizing serositis, hepatitis, interstitial pneumonia, splenitis, orchitis, myocarditis, choroiditis, and enteritis. Gastric nematodiasis. |
| 11 | Male | Adult | Napa Valley, Napa County | *Listeria monocytogenes* encephalitis | Severe necrotizing and histiocytic meningoencephalitis, ependymitis, and choroiditis with gram-positive rods. Multifocal glial scars. Cerebral *Toxoplasma gondii* cysts. Ulcerative keratitis, conjunctivitis with mix-shaped bacteria and yeasts. Severe pleocellular colitis and crypt necrosis. Mild focal granulomatous typhlitis with visceral larva migrans. |
| 12 | Female | Juvenile | Tanzanite Park, Sacramento County | *Staphylococcus aureus* bronchopneumonia | Severe multifocal to coalescing subacute necropsuppurative bronchopneumonia and fibrinosuppurative pleuritis with gram-positive bacterial colonies |
| 13 | Male | Adult | Pismo State beach, San Luis Obispo County | Bacterial bronchopneumonia | Severe multifocal to coalescing suppurative bronchopneumonia with gram-negative rods, gram-positive cocci, and rare foreign material. Minimal multifocal lymphoplasmacytic tracheitis. Moderate multifocal granulomatous lymphadenitis with mineralization and bacteria. Cecal trematodiasis |
| 14 | Male | Adult | Napa Valley, Napa County | Bacterial encephalitis | Moderate multifocal suppurative meningoencephalitis and fibrinosuppurative choroiditis with Gram-negative coccobacilli. Focal glial scar and mineralization. Moderate multifocal neutrophilic and histiocytic pneumonia with foreign material (plant) and bacteria. Mild focal granulomatous colitis with visceral larva migrans |
| 15 | Male | Adult | Oakley, Contra Costa County | Bacterial myofascitis | Skeletal muscle, fascia, salivary gland, and adipose tissue (Pectoral region): Severe diffuse chronic-active fibrinonecrotizing myofascitis, cellulitis, and sialoadenitis with fibrinoid vasculitis, thrombi, and gram-negative bacterial colonies. Severe diffuse edema with thrombi and embolic interstitial pneumonia and fibrinous pleuritis. Heart (right atrium): Thrombus. Cecal trematodiasis. |
| 16 | Female | Adult | Sacramento County | Squamous cell carcinoma | Skin: Squamous cell carcinoma. Severe diffuse acute neutrophilic endometritis with bacteria. Castor gland: Severe diffuse neutrophilic and lymphoplasmacytic adenitis with squamous metaplasia. |
| 17 | Male | Adult | Courtland, Sacramento County | Trauma | Multiple and complete pelvic fractures, myofascial hemorrhages, and edema. Subcutaneous hemorrhages and edema in the head and inguinal region. Mild granulomatous hepatitis with mineralization and Capillaria sp. *Sarcocystis* sp. in tongue. Gastric nematodiasis. |
| 18 | Male | Adult | Sutter County | Capture cardiomyopathy | The pericardial sac contains fibrin and hemorrhage. Moderate multifocal monophasic myocardial necrosis and degeneration. Moderate multifocal glial "track" scars with mild adjacent non-suppurative encephalitis. Mild multifocal granulocytic and histiocytic pneumonia |

Neurological and ocular clinical signs such as circling, ataxia, seizures, blindness, and proptosis were the main clinical signs described in five individuals (S1 Table). Complete animal information including their clinical history and ancillary tests is in S1 Table.

### Mortality and pathological features

The cause of death or the reason for euthanasia in all the studied beavers are in Table 1, which were classified as baylisascariasis, bacterial infections, and non-infectious causes corresponding to 9, 6, and 3 animals, respectively. The latter two categories are subclassified by specific etiological causes below.

### 1. Baylisascariasis

Nine of the 18 beavers (50%) died from lesions associated with neural and visceral larva migrans (NLM and VLM, respectively) consistent with *Baylisascaris* spp. nematodes. Grossly, one beaver showed multifocal to coalescing tan-yellow, <1 mm in diameter, semi-firm granules throughout the cerebrum, cerebellum, and brainstem (Fig 2A). Histologically, this was the most severe case with multifocal eosinophilic granulomas containing a large necrotic core (Fig 2B). In three cases, cavitation tracks surrounded by reactive astrocytes (Fig 2C), foci of gliosis, lymphoplasmacytic infiltration, lymphocytic perivascular cuffing, axonal degeneration (Fig 2D), and scattered eosinophils surrounding NLM (Fig 2E) were seen in different brain sections. NLM within these areas of inflammation was seen in 4 out of the 9 beavers. The most common *Baylisascaris* larva-associated lesion (5/9) in the central nervous system (CNS) was glial scars forming tracks with evident prolonged astrocytic processes (Fig 2F). The latter lesion was also observed in three beavers with different causes of death (Table 2). The most common organ affected by *Baylisascaris* larva migrans (LM) was the brain followed by the gastrointestinal tract (5), lung (4), heart (4), and liver (3). The complete set of organs and tissues affected by *Baylisascaris* LM is in Table 2. The most common brain locations affected by LM were the cerebellum and brainstem. PCR using previously published protocols that have been validated for *Baylisascaris* spp. (Camp et al, 2018) and applied to DNA extracted from FFPE scrolls or frozen brain samples did not amplify products consistent with the *Baylisascaris* positive control. Sanger sequencing on the DNA products indicated non-specific amplification.

**Figure 2.**
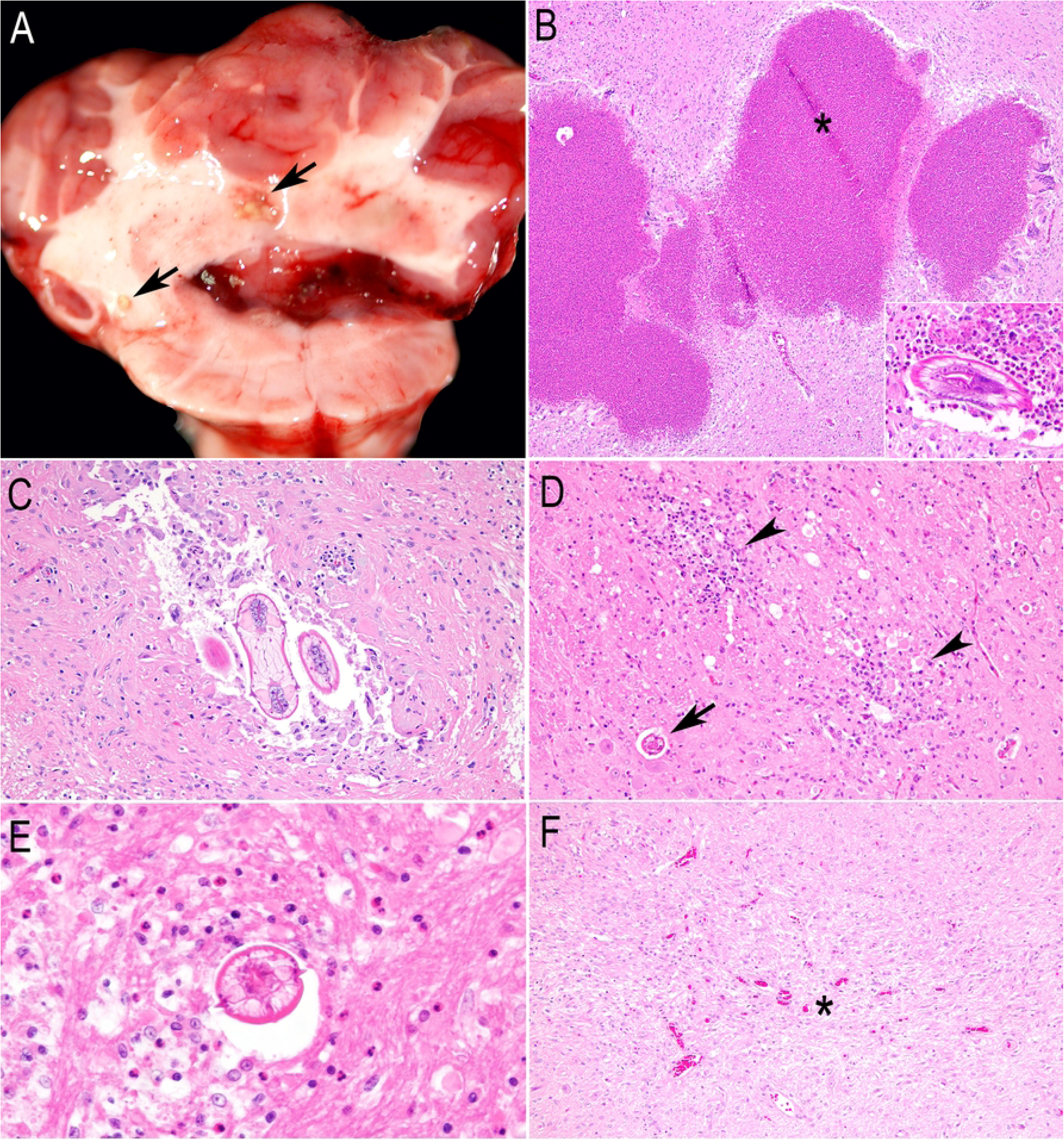
*Baylisascaris* spp.-associated lesions in the central nervous system of beavers in California. A) Multifocal to coalescing tan-yellow granules within the nucleus of the cerebellum compatible with eosinophilic granulomas and prominent congestion/hyperemia of the choroid plexus. B) Microphotograph of Figure 2A showing multifocal eosinophilic granulomas with large cores of necrotic debris (asterisk). Inset: Cross-section of the neural larva migrans (NLM) surrounded by eosinophils. C) Two cross-sections of the NLM of *Baylisascaris* spp. within a cavitation track filled with some macrophages and multinucleated giant cells. The adjacent neuroparenchyma has a dense population of reactive astrocytes and rare lymphocytes. D) Two foci of gliosis, lymphocytes, and axons with dilated myelin sheath and spheroids (arrowheads), and a neuronal larva migrans without evident cellular infiltration (arrow). E) High magnification of the cross-section of a *Baylisascaris* spp. larva with typical lateral alae, excretory columns, and multinucleate intestinal cells surrounded by numerous eosinophils and axonal degeneration. F) Glial scar forming a “migration track” of a larva migrans. There is rarefaction of the neuropil with numerous astrocytes with prolonged processes and neovascularization (asterisk).

**Table 2.**
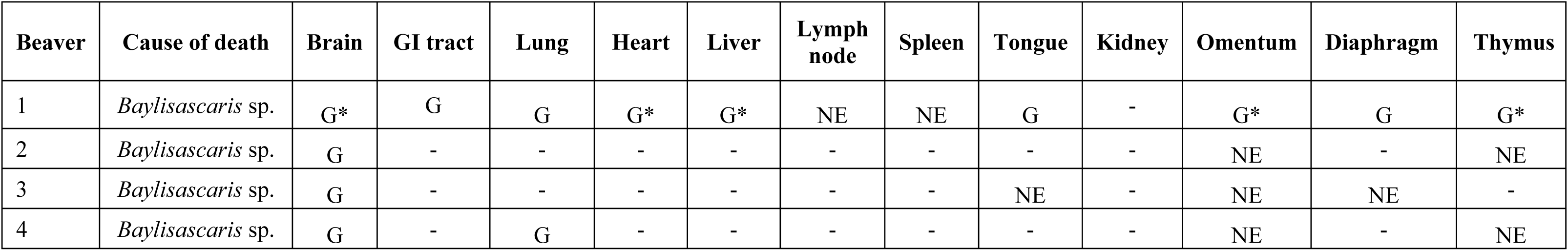
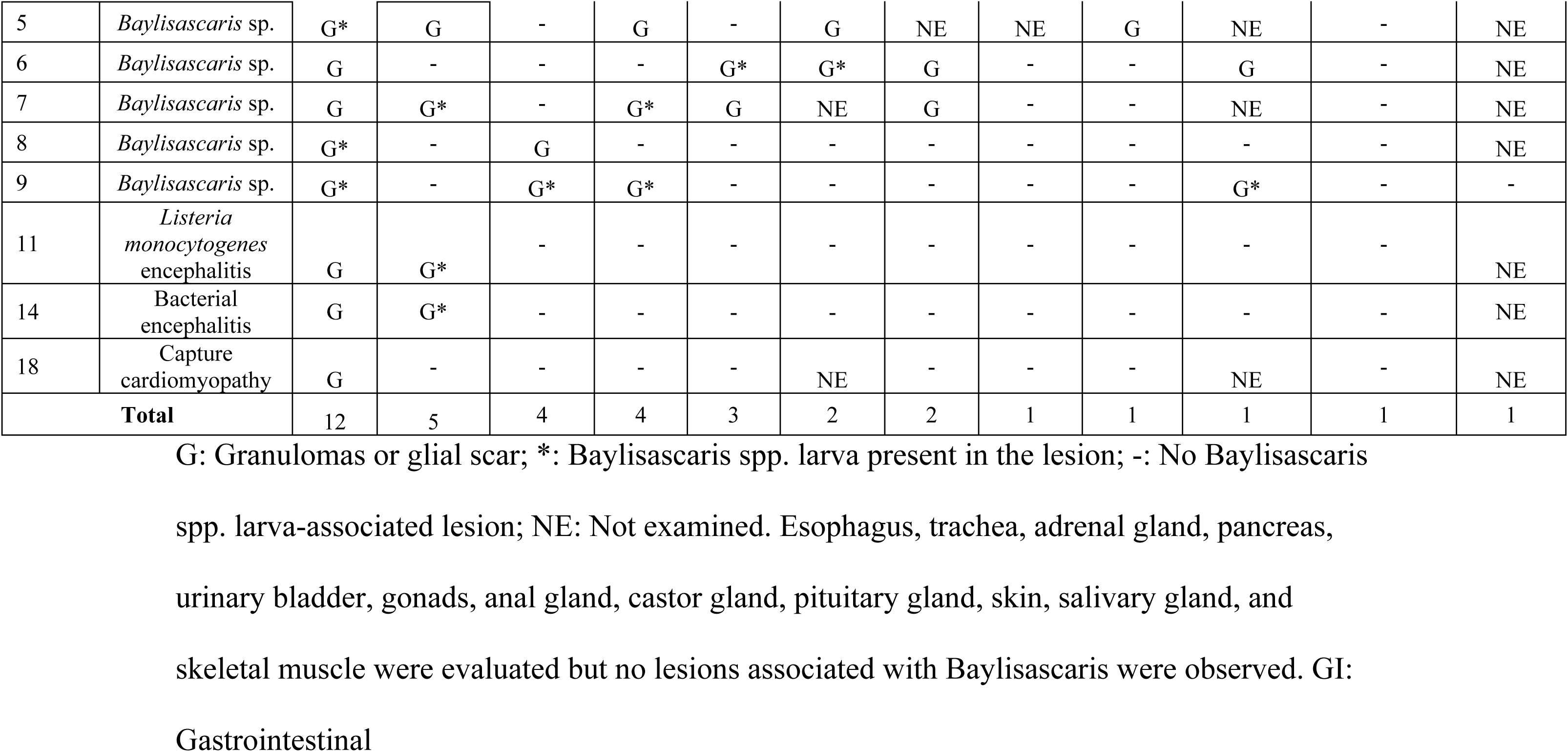
Information on the beavers and their tissues affected by Baylisascaris spp. larva migrans.

|  |  |  |  |  |  |  |  |  |  |  |  |  |  |
| --- | --- | --- | --- | --- | --- | --- | --- | --- | --- | --- | --- | --- | --- |
| 5 | *Baylisascaris* sp. | G* | G | - | G | - | G | NE | NE | G | NE | - | NE |
| 6 | *Baylisascaris* sp. | G | - | - | - | G* | G* | G | - | - | G | - | NE |
| 7 | *Baylisascaris* sp. | G | G* | - | G* | G | NE | G | - | - | NE | - | NE |
| 8 | *Baylisascaris* sp. | G* | - | G | - | - | - | - | - | - | - | - | NE |
| 9 | *Baylisascaris* sp. | G* | - | G* | G* | - | - | - | - | - | G* | - | - |
| 11 | *Listeria monocytogenes* encephalitis | G | G* | - | - | - | - | - | - | - | - | - | NE |
| 14 | Bacterial encephalitis | G | G* | - | - | - | - | - | - | - | - | - | NE |
| 18 | Capture cardiomyopathy | G | - | - | - | - | NE | - | - | - | NE | - | NE |
| **Total** |  | 12 | 5 | 4 | 4 | 3 | 2 | 2 | 1 | 1 | 1 | 1 | 1 |

G: Granulomas or glial scar; \*: *Baylisascaris* spp. larva present in the lesion; -: No *Baylisascaris* spp. larva-associated lesion; NE: Not examined. Esophagus, trachea, adrenal gland, pancreas, urinary bladder, gonads, anal gland, castor gland, pituitary gland, skin, salivary gland, and skeletal muscle were evaluated but no lesions associated with *Baylisascaris* were observed. GI: Gastrointestinal

### 2. Bacterial infections

Six out of 18 beavers died from or were euthanized due to bacterial infections that were systemic or affected the respiratory tract, brain, and musculoskeletal system (Table 1).

#### 2.1 Tularemia

One adult beaver from Lake Tahoe, El Dorado County died of acute systemic random necrotic foci. *Francisella tularensis* was cultured by the CDPH and immunohistochemistry (Fig 3A).

**Figure 3.**
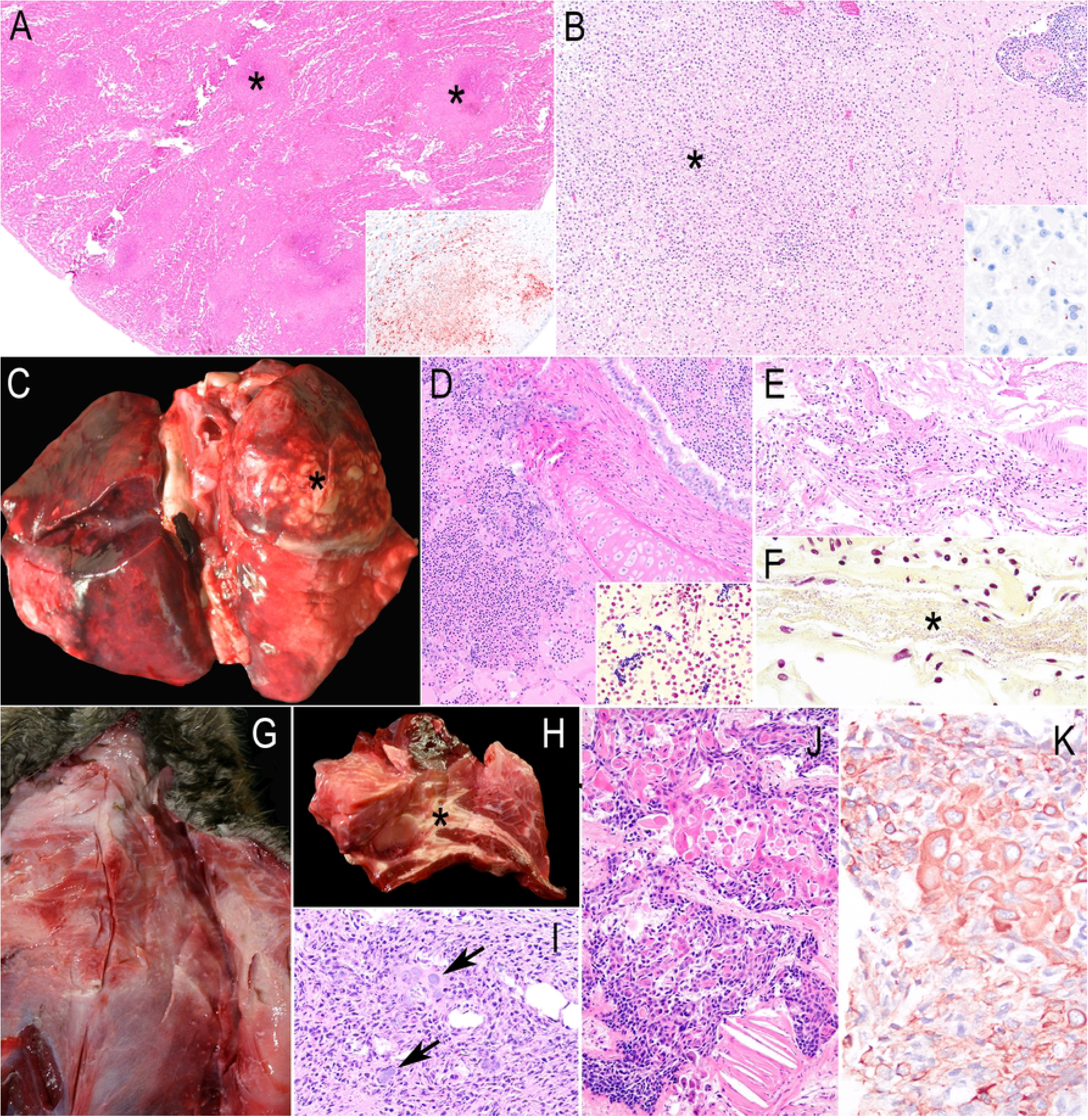
Pathology of the other causes of death besides baylisascariasis in beavers from California. A) Random lytic necrosis in the hepatic parenchyma (asterisks) caused by *Francisella tularensis*. Hematoxylin and eosin (H&E). Inset: Positive immunolabeling for *F. tularensis* by immunohistochemistry in a necrotic focus. Immunohistochemistry (IHC) for *F. tularensis*. B) A focally extensive area of histiocytic and neutrophilic infiltration (asterisk) and severe perivascular cuffing (upper right). H&E. Inset: Positive immunolabeling for intracellular rod-shaped bacteria compatible with *Listeria monocytogenes*. IHC for *L. monocytogenes.* C) Dorsal view of the lungs with severe bilateral bronchopneumonia with evident purulent exudate in the right cranial lobe (asterisk). D) Microphotograph of Figure 3C, numerous neutrophils and necrotic debris are within the bronchus and alveoli. H&E. Inset: Numerous gram-positive cocci among the cellular exudate. Gram stain. E) The leptomeninges are markedly expanded by neutrophils and fibrin. H&E. F) Numerous gram-negative coccobacilli corresponding to *Acinetobacter towneri* are embedded in fibrin strands. Gram stain. G) Ventral view of the thorax, the pectoral muscles are interlaced with tan-white, opaque tissue corresponding to a chronic active inflammation. H) Cross section of the pectoral muscles with evident necrotic exudate in the fascia (asterisk). I) Microphotograph of Figure 3H with coccobacilli bacterial colonies (arrows) surrounded by fibrin and neutrophils among the adipocytes. H&E. J) Squamous cell carcinoma moderate pleomorphic squamous cells with a focal cholesterol granuloma. H&E. K) Positive cytoplasmic immunolabeling of Pancytokeratin (PanCK) in the neoplastic cells shown in Fig 3J. IHC for PanCK.

#### 2.2 Listeria monocytogenes encephalitis

An adult beaver swimming in circles and with ataxia had encephalitic listeriosis with gram-positive rods that immunolabeled for *Listeria monocytogenes* (Fig 3B). This bacterium was cultured from the brainstem and meninges.

#### 2.3 Staphylococcus aureus bronchopneumonia

A beaver affected by a diesel spill was lethargic and hypothermic. Severe bronchopneumonia, primarily affecting the right cranial lung lobe was seen (Fig 3C). Microscopically, neutrophils, fibrin, and necrosis with gram-positive cocci were observed (Fig 3D). *Staphylococcus aureus* was cultured from the lungs, pleura, and liver.

#### 2.4 Bacterial bronchopneumonia

A beaver was found dead with severe bilateral suppurative bronchopneumonia from which *Rahnella aquatilis*, *Streptococcus uberis*, and *Hafnia alvei* were recovered.

#### 2.5 Bacterial encephalitis

A beaver swimming in circles in a river died after showing hypothermia and respiratory distress. Fibrinosuppurative meningoencephalitis and choroiditis with gram-negative coccobacilli (Fig 3E and F) were seen. *Acinetobacter townerii* was cultured from the meninges and cerebrum.

#### 2.6 Bacterial myofascitis

One beaver was found agonal and died. Grossly, coalescing areas of necrosis and granulation tissues were in the left side of the pectoral and cervical muscles, fascia, salivary gland, and adipose tissue (Fig 3G and 3H). Microscopically, necrosuppurative myofascitis and gram-negative bacterial colonies (Fig 3I) were observed. *Aeromonas bestiarum* was isolated from the pectoral and cervical muscles, lung, and liver while *Pasteurella multocida* was cultured from the pectoral muscles.

### 3. Non-infectious causes

#### 3.1 Cutaneous squamous cell carcinoma

A nodule in the right flank was diagnosed as squamous cell carcinoma based on the anastomosed epidermal projections into the dermis, proliferation of atypical squamous cells (Fig 3J). The cytoplasm of the neoplastic cells immunolabeled for PanCK (Fig 3K).

#### 3.2 Trauma

A beaver was found prostrated between two roads in Sacramento and euthanized. Radiographs and gross examination determined comminuted pelvic fractures with myofascial hemorrhages in the lumbar and inguinal regions.

#### 3.3 Capture cardiomyopathy

This beaver was captured, transported, and anesthetized with intravenous anesthetic drugs. Recovery from anesthesia was prolonged. Two days after the procedure, the animal exhibited ataxia, labored drooling, and respiratory distress. Due to the clinical signs, it was euthanized. Grossly, the pericardial sac contained fibrin and hemorrhage. Monophasic myocardial necrosis and degeneration were observed.

### Comorbidities

Parasitic infection was the most common comorbidity. Gastric nematodiasis (Fig 4A) and cecal trematodiasis (Fig 4B) compatible with *Travassosius americanus* and *Stichorchis sp.* were detected in three and four beavers, respectively. Multifocal granulomas with numerous oval embryonated eggs with thick walls and bipolar plugs were seen in the liver (Fig 4C). PCR targeting the 28S rDNA locus yielded a ∼700 bp product that shared 73% identity with *Capillaria plica* (GenBank accession KF836607.1). The beaver diagnosed with listerial encephalitis had rare protozoan cysts in the brain diagnosed as *Toxoplasma gondii* by IHC (Fig 4D), PCR, and sequencing at the ITS1 locus (100% identity). Molecular characterization at the B1 locus demonstrated a Type X variant that contained a previously undescribed single nucleotide polymorphism when compared with other Type X strains that have been described in California and associated with fatal encephalitis in southern sea otters (Shapiro et al, 2019). The cerebral *T. gondii* cysts were not accompanied by any inflammatory cells. *Giardia* spp. cysts were detected in feces from two beavers that died of bacterial encephalitis and capture cardiomyopathy (S1 Table).

**Figure 4.**
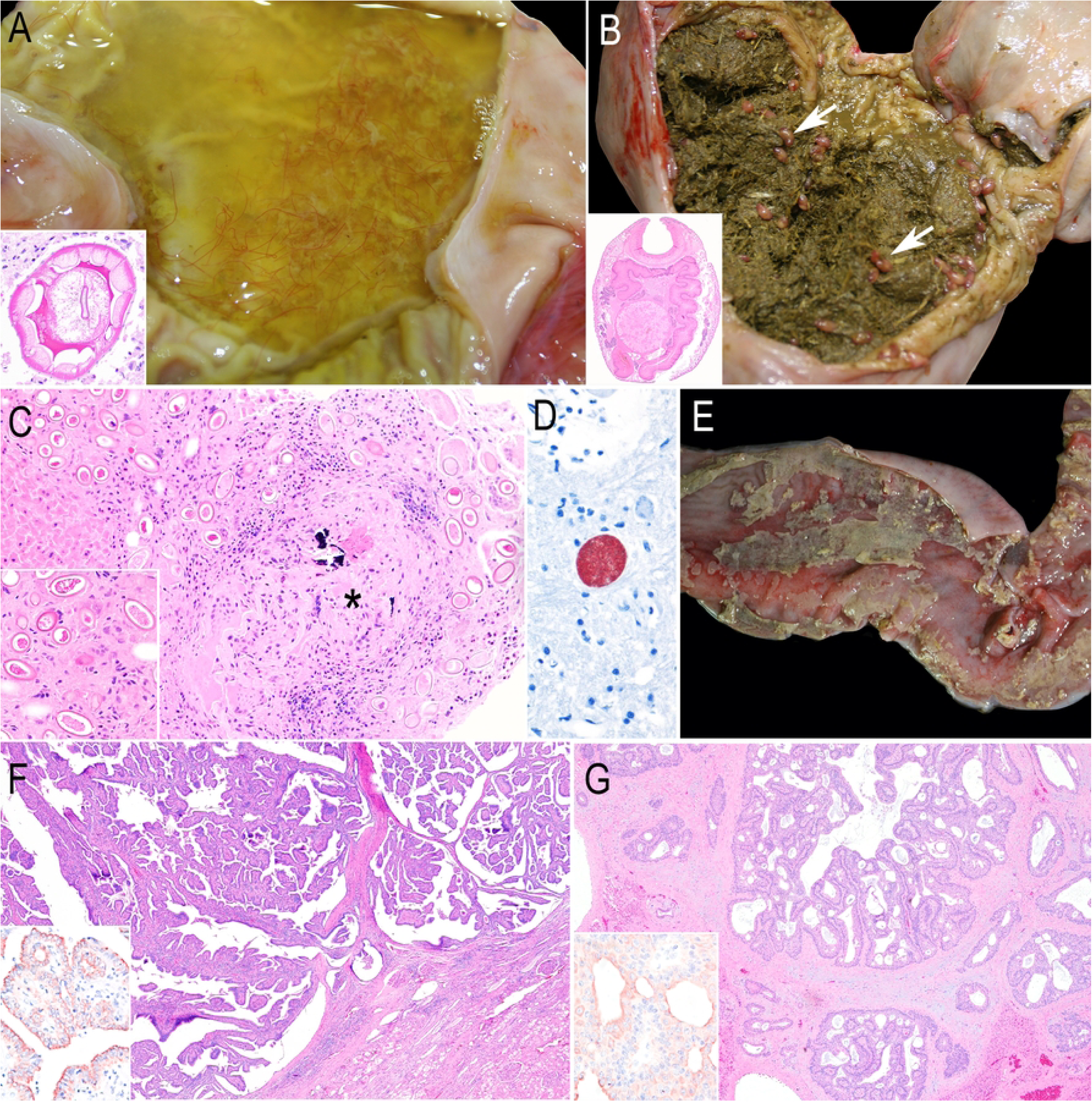
Other comorbidities in free-ranging beavers from California. A) Stomach containing abundant straw-like color, watery fluid with numerous filiform, brown nematodes *Travassosius americanus*. Inset: Cross section of the nematode’s cuticular ridges, platymyarian muscles, lateral cords, and a pseudocoelom with the intestine characteristics of a trichostrongyle. H&E. B) The cecum of a beaver with roughage digesta and numerous trematodes (white arrows) *Stichorchis* sp. Inset: A coronal section of this trematode with oral and ventral suckers and bilateral ceca. H&E. C) A focal granuloma with minimal mineralization (asterisk) surrounded by numerous nematode eggs compatible with a capillarid species. H&E. Inset: The nematode eggs are embryonated and have thick shells and bipolar plugs. H&E. D) Immunolabeling of a *Toxoplasma gondii* cyst in the brain containing numerous zoites. IHC for *T. gondii*. E) Colon of a beaver with diphtheritic membranes attached to the hyperemic mucosa. F) Renal adenoma showing a typical papillary feature. H&E. Inset: The membrane of the cytoplasm of the neoplastic cells is immunolabeled for PanCK. IHC for PanCK. G) Cholangiocellular adenoma dissected by a dense fibrous connective tissue. H&E. Inset: The membranes and occasionally the cytoplasm of a few neoplastic cholangiocytes are immunolabeled for PanCK. IHC for PanCK.

Bacterial colitis (Fig 4E) was diagnosed in two beavers. *E. coli* and mixed flora were recovered from the colon of two beavers while the anaerobes *Bacteroides thetaiotaomicron* and *Terrisporobacter* sp. were recovered in one each.

Two benign tumors, a renal papillary adenoma and cholangiocellular adenoma, were diagnosed in one adult bever. They were solitary, tan-white, and firm nodules. Both tumors were immunolabeled for PanCK and were negative for vimentin (Fig 4F and 4G, respectively).

## Discussion

Approximately 61% (11/18) of beavers submitted to the laboratory were from two contiguous counties in Northern California, Contra Costa and Sacramento. One of California’s main beaver conservation groups is in Martinez (Contra Costa County) [26], which monitors and protects a beaver population. Sacramento County is a highly populated area where human observation of sick beavers is very likely. According to Stallknecht [27], diagnostic submissions and wildlife disease surveillance generally rely on the public’s initial and opportune detection of sick individuals.

Baylisascariasis was diagnosed based on histological features and definitive host distribution. The diagnosis of *Baylisascaris* spp. larval migration might be made by the histological morphology of the larvae, the features of encephalitis, and the presence of definitive hosts in the range of the affected animal [16,28]. Thus, though this study diagnosed baylisascariasis in some beavers, the particular *Baylisascaris* species could not be determined by molecular methods.

*Baylisascaris* spp.-associated lesions were the primary cause of death/reason for euthanasia. Even though multiple adult *Baylisascaris* spp. occur in many wild mammals as definitive hosts [16,25,29,30,31], *B. procyonis*, *B. melis*, and *B. columnaris* found in raccoons, badgers, and skunks, respectively are the most pathogenic members [29]. *B. procyonis* is the archetypal species that produces NLM and has been found in over 150 species of birds and mammals, including humans [30]. Visceral and ocular larva migrans (VLM and OLM, respectively) are also frequently reported [30,31]. NLM of *B. procyonis* was morphologically and molecularly confirmed to cause eosinophilic granulomas in the brains of two beavers from a zoo in Georgia, USA [32]. In free-ranging beavers from Kansas, USA, VLM of *Baylisascaris* spp. was reported to cause eosinophilic granulomas in the liver [33] while a retrospective study in Ontario, Canada revealed encephalitis caused by NLM as the most frequent specific diagnosis in lagomorphs and rodents, including beavers [16]. Similarly, our study reaffirms that beavers can be highly susceptible to *Baylisascaris* spp. NLM and VLM in California. Interestingly, *Baylisascaris* spp.-associated lesions were seen in three animals with different causes of death, suggesting that larval migration throughout the brain is not necessarily fatal, but might leave residual deficits [31]. Possible reasons for the lack of fatal CNS lesions are the low number of eggs ingested, the site and extent of larval migration in the CNS, and the size of the host brain [29,31].

*Baylisascaris* spp. pathogenesis in beavers might be influenced by environmental, parasitic, and host factors. Skunks and raccoons, the definitive hosts of the two most pathogenic *Baylisascaris* spp. in North America, usually defecate in communal sites called latrines, which might thus contain a high load of *Baylisascaris* eggs [29,31,34]. Contaminated feces can be washed away by rainfall into freshwater ecosystems [31] where beavers can ingest larvated eggs and become infected. Ingested eggs hatch in the intestine, penetrate the gastrointestinal mucosa, migrate through the portal circulation to the liver, and along vascular channels to the lung and heart [29,31,35]. In our study, besides the brain, the GI tract, lung, heart, and liver were the most common organs affected by the larval migration. Upon widespread distribution, the larvae migrate to the CNS causing eosinophilic meningoencephalitis, eosinophilic granuloma, and/or “track-like” glial scars [31]. The occurrence of neurological signs is suggested to be related to the number of eggs ingested by the intermediate host [29]. Thus, since all the studied beavers with *Baylisascaris* larval migration had brain lesions, we suspect that the animals might have ingested a large number of infected eggs. Beaver dams decrease the water stream in rivers and freshwater communities, thereby potentially facilitating the accumulation of infective forms of metazoans and protozoans in the ponds [36,37]. Thus, dams built by beavers might increase the exposure to infective eggs of *Baylisascaris* spp.

Six beavers died of different bacterial infections. Tularemia, a zoonotic bacterial disease caused by *F. tularensis*, was diagnosed in one beaver in Lake Tahoe, CA. In North America, lagomorphs such as snowshoe hares (*Lepus americanus*) and rodents such as muskrats (*Ondatra zibethicus*) and beavers are most reported by *F. tularensis*. Arthropods such as ticks play a significant role in the transmission [11,38]. Beavers and other rodents are predominantly involved in the aquatic cycle of tularemia, which is associated with the subspecies *F. tularensis holarctica* [11,39]. This subspecies can persist for long periods in water courses, ponds, lakes, streams, and rivers [38]. Lake Tahoe is a popular recreational area [40]. Thus, the presence of *F. tularensis* raises concerns about the contact between humans, wildlife hosts, and vectors of this zoonotic bacterium.

*Listeria monocytogenes* has been isolated from rectal swabs of beavers as well as other wild mammals, birds, and reptiles. Some of these were hypervirulent strains that could be fatal in humans and domestic animals [41]. The *L. monocytogenes* encephalitis case shared features with fatal neurolisteriosis caused by hypervirulent genotypes in domestic ruminants [42]. The fact that this bacterium can persist in the soil and water and can be shed by wild and domestic animals suggests that beavers and other inhabitants of freshwater ecosystems might be at risk for listeriosis [42,43].

Fatal bacterial bronchopneumonia was diagnosed in two beavers. One died of *Staphylococcus aureus* infection and the other one by a combination of gram-negative and positive bacteria (*Rahnella aquatilis*, *Streptococcus uberis*, and *Hafnia alvei*). *S. aureus* seems to be part of the normal conjunctival microbiota of North American beavers [44] and was isolated from the nasal cavity of beavers (exact species was not recorded) [45]. *S. aureus* was recovered from multiorgan abscesses, pyelonephritis, lymphadenitis, pyuria, and pneumonia from Eurasian beavers (*Castor fiber*) [46]. One studied beaver suffered the stressors of an oil spill-over event suggesting that *S. aureus* might be a fatal opportunistic pathogen. In the other case, *Hafnia alvei, Rahnella aquatillis*, and *S. uberis* are water-borne pathogens that in rare instances cause sepsis or purulent inflammation in humans [47,48,49]. The latter bacterium is also a major agent in bovine mastitis and dairy cow environments [50].

Bacterial encephalitis and myofascitis caused by *Acinetobacter towneri* and *Aeromonas bestiarum*, respectively, were diagnosed in one beaver each. The latter was recovered from other organs suggesting septicemia. *A. towneri* is a gram-negative bacterium that has been recovered from activated sludge [51] and swine feces [52]. Even though this species has not been associated with clinical cases, other *Acinetobacter* species, such as *A. baumannii*, are potential nosocomial pathogens causing pneumonia, urinary tract infections, and meningitis in hospitals [53]. *A. bestiarum* is found in aquatic environments causing septicemia or skin ulcers in carps [54]. In mammals, it was recovered from a red squirrel (*Sciurus vulgaris*) [55]. To the authors’ knowledge, no reports of the bacteria described above have been previously documented in North American beavers.

Regarding the non-infectious causes, cutaneous squamous cell carcinoma (SCC) was diagnosed in one beaver. Even though tumors in beavers are rarely documented [56], SCC arising from the esophagus in a captive beaver has been documented[57]. The potential causes of this SCC are unknown. In the beaver with pelvic fracture, the clinical suspicion was trauma by vehicle collision. A large-scale study about traumatic injuries caused by motor vehicle collisions in wildlife found that the abdomen and pelvis were the most frequently affected body segments; specifically in rodents, the most common trauma was hip fractures [58]. Car accidents have been reported as a plausible cause of death in beavers [59]. Capture, restraint, transport, and anesthesia in free-ranging animals are necessary for translocation, sample collection, rescues, etc. However, stress during these procedures has the potential to cause cardiomyopathies and capture myopathy [60,61]. Timing, recovery periods, and mitigating sources of stress need to be considered. In the beaver with capture cardiomyopathy, contraction band necrosis was a predominant feature, and this is typically associated with stress [60,62]. The fact that the heart and not the skeletal muscle was primarily affected in this beaver might represent the acute course of the syndrome [60].

The most frequent comorbidity in the studied beavers was endoparasitism. Gastric trichostrongyles and cecal trematodes were morphologically identified as *Travassosius americanus* [15,63] and *Stichorchis* sp. [63,64], respectively. These are species that commonly infect beavers without eliciting significant disease. Granulomatous hepatitis with numerous eggs compatible with capillarid nematodes was observed [65]. Within this group, *Capillaria hepatica* has a worldwide distribution and typically infects the liver of numerous species of rodents including muskrats and beavers [66], lagomorphs, monkeys, humans, and other mammal species [67,68]. In the majority of species, the pathogenicity of *C. hepatica* in the liver of rodents is low [68] as was observed in the studied beaver. The possible diagnosis of *Capillaria* sp. suggests the presence of embryonated eggs in the aquatic environment representing a potential zoonosis. Other potential zoonoses identified in the study population were *Toxoplasma gondii* and *Giardia* spp. *T. gondii* was previously identified in free-ranging [69] and captive [70] beavers. The strain identified in the studied beaver was a type X variant not previously described but nearly identical to Type X and other Type X variants that appear to be virulent in Southern sea otters [24]. In the beaver in this study, the parasite tissue cysts did not elicit evident inflammation in the brain.

Further studies should be conducted to evaluate the susceptibility of other animal species for fatal infections by this *T. gondii* strain. *Giardia* spp. was previously reported in beavers from Canada and the USA [71,72,73]. Beavers are considered amplification hosts of *Giardia* spp. that can contaminate freshwater ecosystems, thereby contributing to the risk of giardiasis in humans [73].

Two beavers had bacterial colitis by common commensal bacteria of the large intestine of humans, but they can become pathogenic in immunocompromised individuals [74,75]. Since both beavers had fatal baylisascariasis with depression and neurological signs, the colitis was likely caused by the opportunistic effect of the colonic bacterial microbiota.

Two benign tumors were diagnosed in one adult beaver, a chollangiocellular adenoma and a papillary renal adenoma, which have not been reported in beavers before [76]. Both tumors were considered incidental findings.

## Conclusions

This study shows that free-ranging beavers in California can carry and succumb to numerous infectious diseases considered hazards to domestic animals and humans. Since these species frequently visit the natural freshwater habitats of beavers, we believe that studying the diseases of this rodent might be an excellent barometer to determine the freshwater ecosystem health as with other aquatic animals [77,78]. Because beavers are ecosystem engineers, they have been historically translocated to facilitate the restoration of degraded ecosystems [79]. Our study and others [15,80] alert to the potential risk of moving pathogens that might affect other beaver populations, native wildlife, domestic animals, and humans. Consideration should thus be made when handling beavers and planning translocations to mitigate risks of unintended introduction of infectious disease agents.

## Acknowledgments

We thank the pathologists, faculties, and technicians from the CAHFS, Davis Laboratory for having worked on these cases. Special thanks to CDFW staff, biologists, and personnel of the rehabilitation centers, especially to Lindsay Wildlife Experience for their hard work in understanding and improving beaver health. We also thank Melinda Houtman for creating the map and Lezlie Rueda and Chloe Resngit who supported the molecular parasitology efforts.

## Notes

### Competing Interest Statement

The authors have declared no competing interest.

